# Exponential magnetophoretic gradient for the direct isolation of basophils from whole blood in a microfluidic system

**DOI:** 10.1101/2022.02.11.480005

**Authors:** Nicolas Castaño, Sungu Kim, Adrian M. Martin, Stephen J. Galli, Kari C. Nadeau, Sindy K.Y. Tang

## Abstract

Despite their rarity in peripheral blood, basophils play important roles in allergic disorders and other diseases including sepsis and COVID-19. Existing basophil isolation methods require many manual steps and suffer from significant variability in purity and recovery. We report an integrated basophil isolation device (i-BID) in microfluidics for negative immunomagnetic selection of basophils directly from 100 μL of whole blood within 10 minutes. We use a simulation-driven pipeline to design a magnetic separation module to apply an exponentially increasing magnetic force to capture magnetically tagged non-basophils flowing through a microtubing sandwiched between magnetic flux concentrators sweeping across a Halbach array. The exponential profile captures non-basophils effectively while preventing their excessive initial buildup causing clogging. The i-BID isolates basophils with a mean purity of 93.9%±3.6% and recovery of 95.6%±3.4% without causing basophil degradation or unintentional activation. Our i-BID has the potential to enable basophil-based point-of-care diagnostics such as rapid allergy assessment.

## Introduction

Basophils are the rarest type of granulocytes in peripheral blood, comprising 1% or less of all leukocytes. Despite their low abundance, basophils are important regulatory and effector cells in a wide range of immune functions and disorders (*1*–*4*). In particular, basophils play critical roles in allergic diseases. When triggered by stimulants such as food or environmental allergens, basophils can secrete a range of mediators (e.g., histamine, leukotrienes, platelet activating factor) to induce inflammatory reactions (*5*–*7*). They can infiltrate tissues undergoing atopic hypersensitive inflammation (*8*), and act as initiators of allergic inflammation (*4*–*6*, *9*) and possibly of anaphylactic reactions (*10*–*13*). Basophils have also been implicated in other diseases (e.g., enhancing the innate immune response against sepsis (*14*), producing angiogenic factors in cancer tumorigenesis (*15*), and modulating humoral response against SARS-CoV-2 (*16*)), and in immunoregulatory functions (e.g., imprinting alveolar macrophages (*17*), promoting Th2 cell differentiation (*18*, *19*), and recruiting eosinophils (*20*, *21*)).

The identification and isolation of basophils are important for both clinical and research studies. For example, the activation of basophils in the presence of allergens *ex vivo* has been found to correlate with the allergic status of the subject. The basophil activation test (BAT), an *ex vivo* blood test, has demonstrated high specificity (75-100%) and sensitivity (77-98%) in the diagnosis of allergy to a range of allergens including peanut, cow’s milk, egg, therapeutic drugs, and pollen (*22*–*26*). Here, basophils are typically identified by gating schemes in flow cytometry data, commonly as SSC^low^/HLA-DR^-^/CD123^+^. Basophil activation is then measured by the level of expression of activation markers CD63, CD203c, and/or avidin (*27*–*29*). In fundamental research, bulk and single-cell RNA sequencing of basophils has uncovered mechanistic pathways and transcriptional profiles associated with basophil differentiation (*30*, *31*) and function (*6*, *15*, *17*, *32*, *33*). The accuracy of RNA-sequencing, especially deep transcriptional profiling of specific cell types, relies on starting with purified target cell populations—these are currently most commonly isolated by fluorescence-activated cell sorting (FACS) (*34*–*36*).

Since basophils are rare, their isolation from whole blood has been challenging. Conventional methods of separation are based on density gradient, FACS, immunomagnetic negative selection, or a combination of these methods. Shiono *et al*. demonstrated a flow-through basophil isolation method using 5 Percoll density gradients, but their approach suffered from relatively low recovery (~56-77%) and purity (~51-72%) (*37*). Degenehart *et al*. used FACS to isolate viable basophils from white blood cells (WBCs) enriched from venous blood with a modest mean purity of 84% (range 75-95%) and a low mean recovery of 20% (range 15-30%) (*38*). This method required upstream removal of red blood cells (RBCs) by density gradient or lysis. The need for surface markers may also affect downstream assays that require antibody labeling. The best performing basophil isolation methods reported thus far employ immunomagnetic negative selection of basophils (*39*). Using a commercial immunomagnetic negative selection kit (*40*), basophils were isolated with a mean purity of 99.3% (range 97-100%) from WBCs, where RBCs were first removed using an RBC aggregation agent followed by centrifugation. Despite the high purity, this method has several drawbacks: 1) The recovery was modest, with a mean recovery of 75.6% and a large variability in performance (range 39-100%) (*41*). 2) It required a large volume (30 mL) of blood. 3) The performance of isolation from whole blood was not reported and is expected to be reduced due to the loss of basophils during the RBC removal step. 4) It required many manual pipetting and centrifugation steps which subject leukocytes to shear stresses that can alter their immunophenotype (*42*). Although faster than approaches using multiple Percoll gradients combined with immunomagnetic negative selection (which took 4 hours) (*43*), this process still required ~90 min (*41*, *44*).

Herein, we describe the design and development of an integrated microfluidic basophil isolation device (i-BID) to perform immunomagnetic negative selection of basophils from whole blood. We chose negative selection instead of positive selection to maintain basophils in their native, unlabeled state. The device consists of deterministic lateral displacement (DLD) channels to enrich WBCs from whole blood, a mixer to mix magnetic nanoparticles (MNP) and negative selection antibody (NSAb) with enriched WBCs, and a magnetic separation device (MSD) to deplete non-basophils (Fig. 1A).

**Fig. 1.**
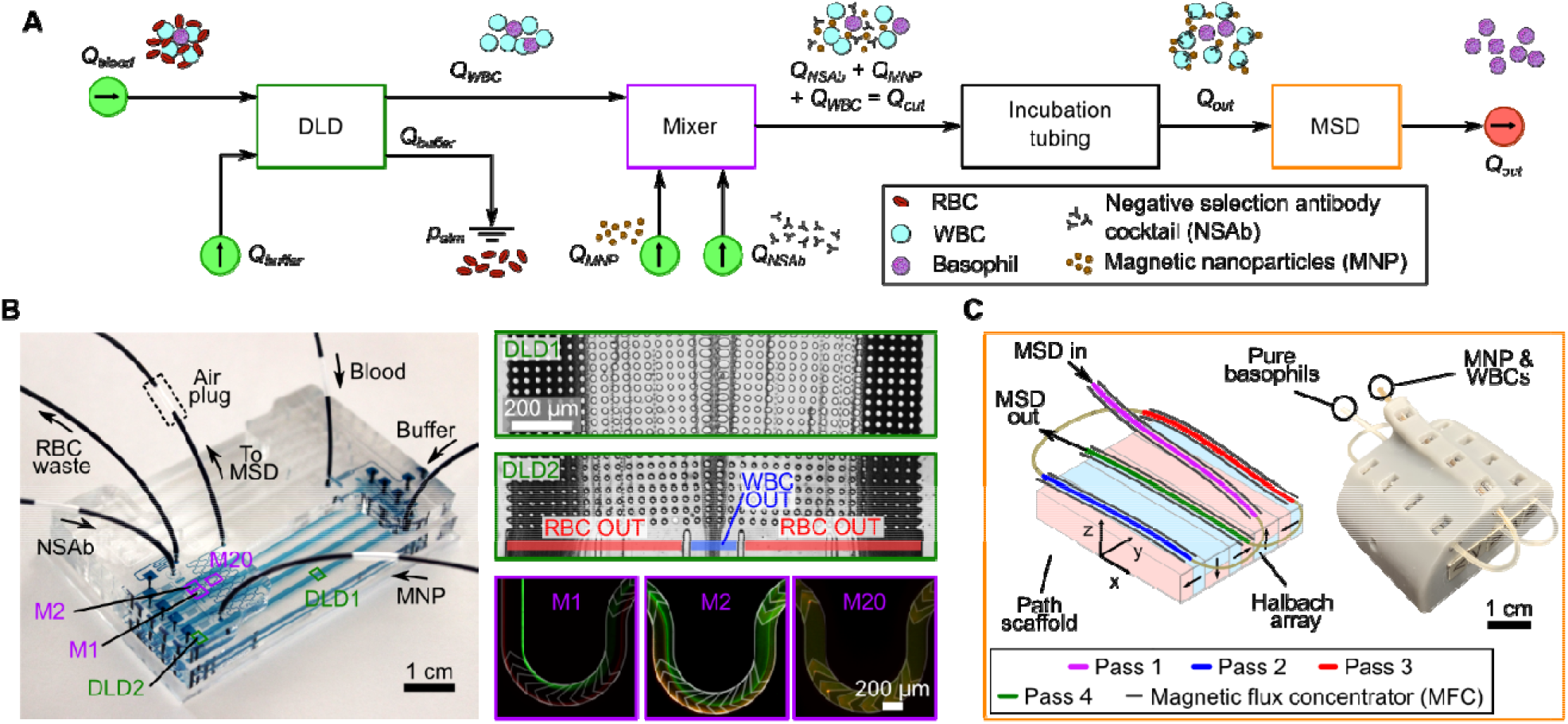
Overview of the basophil isolation device (i-BID) process flow. (A) Schematic diagram of the microfluidic circuit for WBC enrichment and basophil isolation by magnetophoretic negative selection. All injection (green) and withdrawal (red) flow rates were controlled with syringe pumps. (**B**) Image of the microfluidic chip (left). Microscopy images of deterministic lateral displacement (DLD) stages DLD1 and DLD2, and mixer units M1, M2, and M20 are shown in the insets. The center outlet of the DLD channels collects enriched WBCs. The mixer channel mixes two fluorescent streamlines by M20 (right) to demonstrate mixing of WBCs with negative selection antibody (NSAb) and magnetic nanoparticles (MNP). (**C**) Schematic diagram and photograph of the magnetic separation device (MSD). The diagram shows the locations of 4 passes of tubing with steel wires acting as MFCs running parallel. The image shows brown-tinted fluid entering and clear fluid exiting, qualitatively demonstrating the depletion of MNP in the MSD.

The key innovation lies in the utilization of a model-driven design pipeline for the development of the MSD to apply an exponentially increasing magnetophoretic gradient to capture non-basophils gradually, as the cells flow through, on the inner walls of a polyethylene tubing. This approach contrasts with existing bulk or microfluidic magnetic separation methods that have a single region with constant magnetic force or discrete regions of differing magnetic forces (*45*–*47*). A gradual initial increase in magnetophoretic force prevents an abrupt, excessive buildup of captured non-basophils and unbound MNP, which could lead to clogging and non-specific capture of basophils. The steep increase in magnetophoretic force at the end of the MSD serves to capture non-basophils with low numbers of MNP and any unbound MNP. The MSD uses steel wires as magnetic flux concentrators (MFCs) and a 3D printed scaffold to sweep the path of the tubing across an array of magnets, which consists of five magnets arranged in a Halbach configuration. The spacing of the MFC wires and the tubing relative to the Halbach array is determined by a target exponential magnetophoretic gradient profile and numerical simulations of the resultant magnetic fields across the tubing and the MFC. Between each run, a new tubing is inserted into the MSD without the need for fabricating new microchannels or re-aligning the flow path with the magnets. The i-BID achieves a mean purity of 93.9% (range 85.5-98.2%) and a mean recovery of 95.6% (range 91.0-100.0%) from 100 μL of blood, which is too small a volume to process using bulk isolation methods. The entire process takes approximately 10 min from the loading of whole blood to the retrieval of isolated basophils, which remain viable and inactivated.

## Results

### Overall process flow of the integrated basophil isolation device

Fig. 1A and Supplementary Video 1 show the process flow of our integrated basophil isolation device (i-BID). It consists of four stages: 1) DLD channels to enrich WBCs from whole blood, 2) serpentine channels with Herringbone grooves to mix MNP/NSAb with enriched WBCs, 3) a section of tubing whose length determined the incubation time for tagging non-basophils with MNP, 4) the MSD for removing non-basophils from the suspension. Here, we selected DLD channels to separate WBCs from RBCs because of the high purity and recovery of WBCs (see Fig. S1) and the compatibility of its operating flow rates with downstream mixing of MNP/NSAb and basophil isolation in the MSD. Following the DLD, the enriched WBCs were mixed with MNP and NSAb in a mixer consisting of a serpentine channel with Herringbone grooves (*48*). Dean and helical flows induced by the channel’s serpentine bends and Herringbone grooves, respectively, promoted efficient mixing of MNP and NSAb with WBCs and increased the incidence of binding events between cells and MNP. Insets in Fig. 1B shows qualitatively that separate streamlines were well mixed before the exit of the mixer. In transit to the MSD, the MNP/NSAb and WBC were incubated in the incubation tubing for 3 min, which translated to a tubing length of 27 cm when operating at a blood injection flow rate of 3 mL/hr. The incubation tubing was fed to the inlet of the MSD to capture magnetically tagged non-basophils and excess MNP (Fig. 1C). The outlet of the MSD contained enriched basophils, which were collected to assess purity, recovery, and activation status.

### Design and operation of the MSD

The concentration of basophils in peripheral blood is approximately 1% of WBCs, corresponding to ~10 cells/μL of whole blood (*2*). In contrast to the isolation of circulating tumor cells (CTCs) which are present at much lower concentrations (less than a few cells per mL), basophil isolation requires a smaller volume of blood than CTC isolation does. This difference allowed us to design a magnetic-activated cell sorting (MACS) device based on capture mode instead of deflection mode because the magnetically tagged non-basophils and unbound MNP could be sequestered adequately on the inner walls of a microtubing. The capture mode also offered a simple means to integrate basophil isolation with downstream on-chip processes. There is no need to match the fluidic resistance of the product and the waste outlets, which would be necessary in devices operating in deflection mode.

Fig. 1C shows a schematic diagram and a photograph of the MSD consisting of a 3D-printed scaffold housing a Halbach array, a polyethylene microtubing threaded over the Halbach array in four passes, and steel wires acting as magnetic flux concentrators (MFCs) positioned parallel to the tubing on its left and right. The Halbach array comprised 5 magnets arranged in a Halbach configuration. We chose to use a Halbach array because the resulting magnetic field 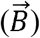 was stronger, more far-reaching, and symmetric about the center magnet than other possible configurations of the 5 magnets. We chose the positions of the tubing and the MFCs to achieve an exponentially increasing magnetophoretic force along the length of the tubing, i.e., the path of the cells, to gradually capture magnetically tagged cells and unbound MNP from the flow. The gradual initial increase in the magnetophoretic force prevented abrupt accumulation of magnetically tagged cells and MNP which could lead to clogging and undesired trapping of basophils. The steep increase in magnetophoretic force at the end of the MSD helped to remove non-basophils with low magnetic susceptibility and ensured that any unbound MNP and magnetic debris did not escape the MSD and contaminate the product of purified basophils. In principle, other magnetophoretic force profiles that are slowly increasing should also work. Here we selected an exponential profile since it was compatible with the Halbach array consisting of 5 magnets and 4 passes across the magnetic field, but the design pipeline can accommodate other target profiles. Four passes were chosen to maximize the length of tubing that could span the 5 magnets while avoiding physical overlap between adjacent passes. Based on the polarity of the magnets in the Halbach array, we chose to sweep the tubing along the long axes of the magnets because it resulted in a more uniform magnetic force along the tubing than to have the tubing traverse across all magnets which would result in abrupt changes in the magnetic force. Since the tubing was sandwiched laterally by the MFCs, positioning the tubing over magnets 1, 3, and 5 (see Fig. 2A), where 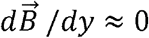, allowed the magnetic flux to be concentrated inside the tubing by the MFCs (see details in Methods).

**Fig. 2.**
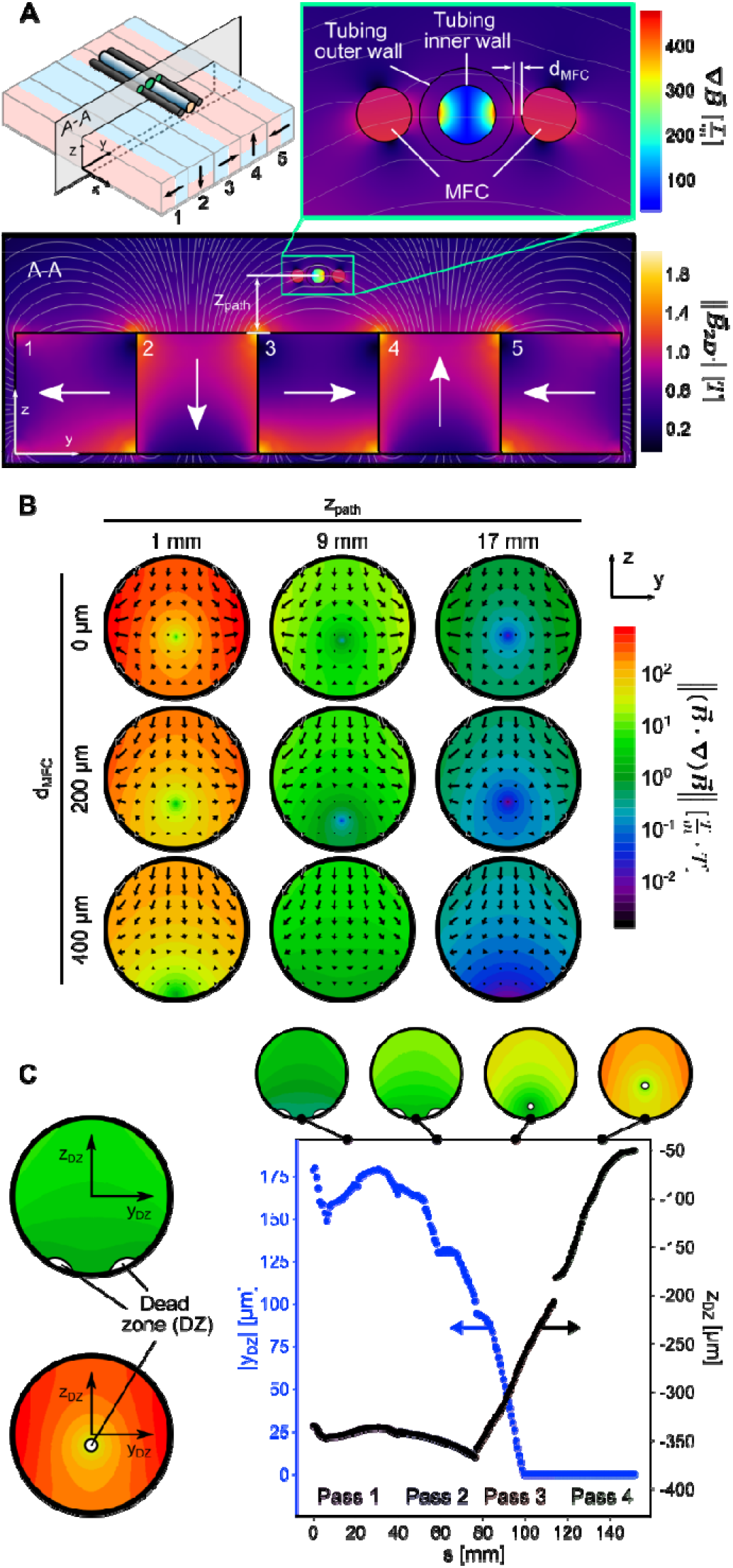
Simulation data used for the design of the magnetic separation device (MSD). (**A**) Representative simulation data from the parametric sweep for z_path_ and d_MFC_ (distance between cell path and magnetic flux concentrators (MFCs)) at a center-plane A-A. The rainbow colormap shows magnetic flux density gradient and the thermal colormap shows the norm of the magnetic flux density. (**B**) The resulting magnetic force field norm in the tubing across different values of z_path_ and d_MFC_. The colormap represents the magnetic force field norm and the arrows indicate the direction of the magnetophoretic force. (**C**) The variation in dead zone (DZ) position, where the net magnetic force field is zero. The origin of (y_DZ_, Z_DZ_) is at the center of the tubing cross section. Inflections in the plot are due to changes in the finite element mesh when the geometry changes as we sweep over different values of z_path_ and d_MFC_. Discontinuities between passes are due to the edges of the magnets when the tubing path turned around to pass over the Halbach array again.

We used numerical simulation in COMSOL (AC/DC module) to model the magnetic field and magnetophoretic forces at different locations relative to the Halbach array (see Table S1 for model parameters). The goal of the simulations was to identify a set of spatial coordinates to position the tubing and the MFCs relative to the magnets to achieve the target exponential magnetophoretic force profile along the path of the tubing. These spatial coordinates were then used to inform a 3D printed model of the MSD scaffold to hold the tubing, the MFCs, and the magnets. To save computation cost, our approach started with a 2D simulation of the magnetic fields in the yz plane (Fig. 2A) where we varied the relative distance between the tubing and the magnets (z_path_), and between the tubing and the MFCs (d_MFC_). After that, we identified combinations of z_path_ and d_MFC_ that could give an exponential magnetophoretic force profile, and then translated these distances to actual 3D coordinates for the tubing and the MFCs.

The governing equation for the magnetophoretic force that a magnetic field exhibits on a magnetic particle is given in Eq. 1 (*45*, *49*),

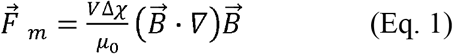

where V is the volume of the particle, Δχ is the difference in magnetic susceptibilities between the particle and the fluid, *μ*_0_ is the magnetic permeability in a vacuum (4 π × 10^-7^ Tm/A), and 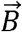 is the magnetic flux density. We designed the MSD around the 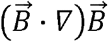 term, known as the magnetic force field (*50*), since the magnetophoretic force scales directly with this term and *V*Δ*χ*/*μ*_0_ is constant for a given size and material of the particles.

To vary 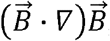, we varied z_path_ and d_MFC_. We performed a parametric sweep to model the magnetic flux concentration by the MFCs as a function of z_path_ and d_MFC_ in 2D at the midplane of the Halbach array above magnet 3 where 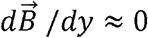 (section A-A in Fig. 2A). Fig. 2B shows the distribution of the magnitude and the direction of the magnetic force field inside the tubing for a few combinations of z_path_ and d_MFC_. As expected, the smallest z_path_ (=1 mm) and d_MFC_ (= 0 μm) yielded the highest maximum magnitude of the magnetic force field within the cross section of the tubing (i.e., 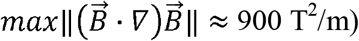), corresponding to the strongest magnetophoretic force. When d_MFC_ was small, 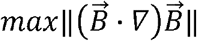 occurred close to the left and the right rims of the tubing. This distribution was expected since the MFCs were positioned to the left and the right of the tubing. Even when d_MFC_ was large, the influence of the MFCs on the magnetic flux was evident in the non-zero horizontal components of the magnetic force field. For most combinations of z_path_ and d_MFC_, both lateral and vertical-pointing forces were present (also see Fig. S2), thereby allowing the MNP-tagged cells to be captured across a greater area of the inner wall of the tubing compared with other MACS devices with strictly lateral or vertical-pointing magnetic forces. In addition, the “dead zone”, i.e., the region with zero net magnetic force field (see Fig. 2C), did not maintain a constant position along the tubing, and therefore, the tagged cells in all streamlines within the tubing would experience a force to be captured.

Fig. 3 shows our process to identify the spatial coordinates (x, y, z) of the entire length of the tubing and the MFCs to achieve an exponential magnetophoretic force profile. Our process consisted of four steps.

**Fig. 3.**
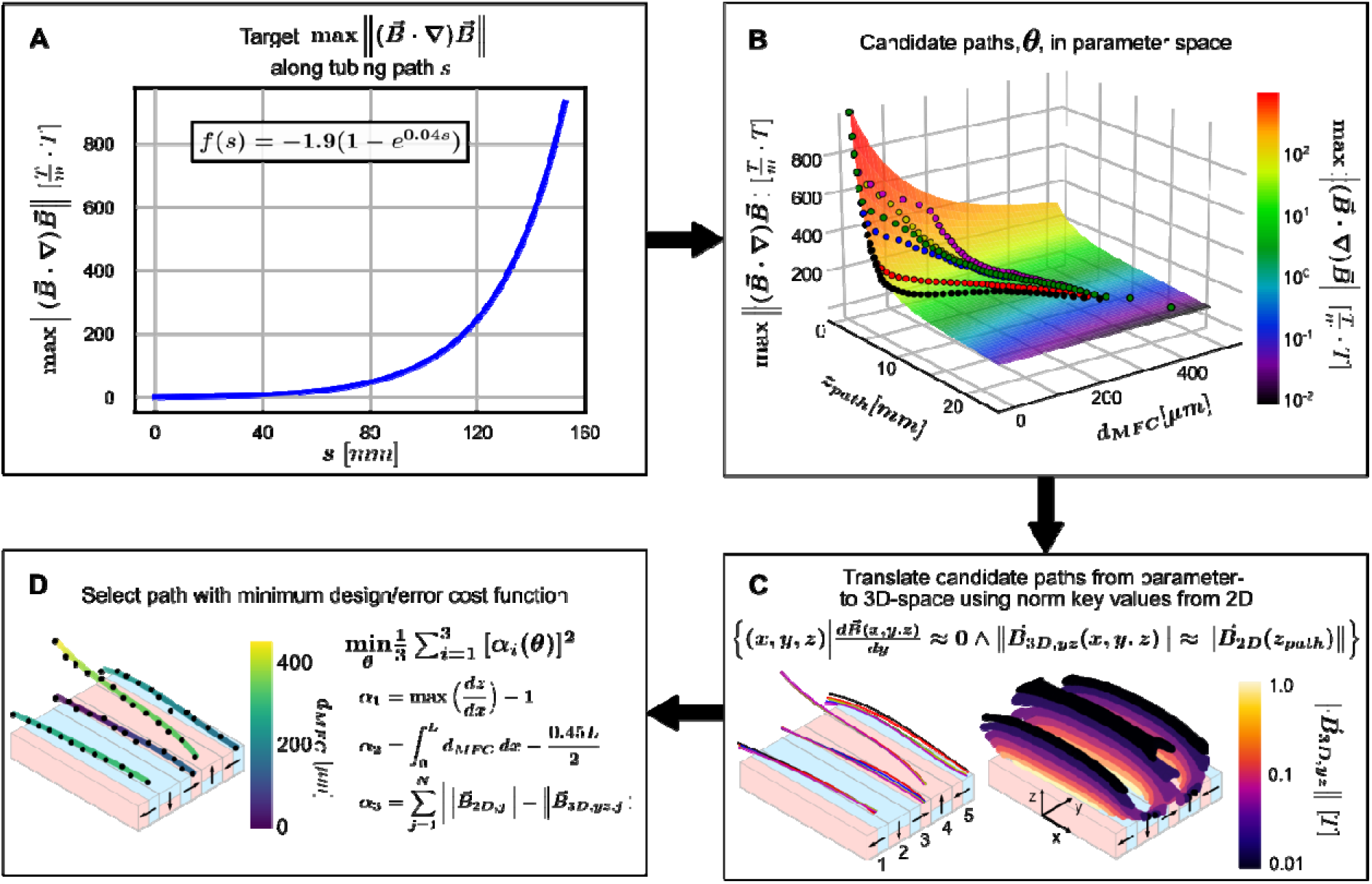
Magnetic separation device (MSD) design pipeline. (**A**) We started with a target magnetophoretic profile across the tubing length which we quantified by the maximum value of in the cross section of the tubing at a given position *s*. (**B**) Using the target profile as a guide, we identified all the paths *θ* that would generate the target profile when ascending the parametric surface which we defined with 2D simulations (see Fig. 2). (**C**) Candidate parametric paths θ were translated to physical 3D candidate paths by finding the point at each position in the tubing path that satisfied the criteria listed in the diagram. (**D**) The best path was selected to minimize sharp changes in the tubing path (*α*_1_), promote gradual changes in d_MFC_(*α*_2_), and minimize error between the 2D and 3D field norms (*α*_3_).

In step 1, we generated a target mathematical profile of 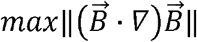 as a function of the path of the tubing *s* (Fig. 3A). The upper bound of the target value for 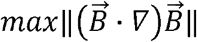 in Fig. 3A was set at 900 T^2^/m, equal to the highest value of 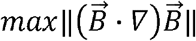 for z_path_ = 1 mm and d_MFC_ = 0 μm in our 2D simulation (see Fig. 2B). We selected an exponential function *F*(*s*) = –1.9(1 – *e*^004*s*^) as a test target profile. Other similar functions are also compatible with our design. The path length *s* spanned the length of 4 passes of the tubing across the long axes of the magnets (152.4 mm). We ignored the portions of the tubing that looped back outside the MSD since they were exposed to weak magnetic flux.

In step 2, we identified different combinations of z_path_ and d_MFC_ that could generate an exponential magnetophoretic force profile along the path of the tubing. To do so, we plotted 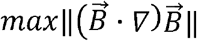 obtained from the parametric sweep (as described in Fig. 2B) on a surface as a function of z_path_ and d_MFC_ (Fig. 3B). We then identified candidate paths (*θ*) on the 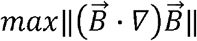 surface that would generate the exponential profile *f*(*s*). Each candidate path consisted of approximately N = 150 points on the 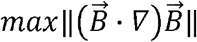 surface. Each point corresponded to one set of z_path_ and d_MFC_ values.

In step 3, we translated the candidate parametric paths into physical paths in 3D (Fig. 3C). The values of d_MFC_ were already defined from the candidate parametric paths in step 2, but z_path_ had to be translated into physical positions in 3D. We performed a high-resolution numerical simulation of the magnetic field in free space in 3D over the Halbach array, and identified the set of (x, y, z) points in the 3D simulation that satisfied the following two criteria: 1) 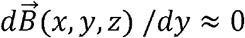, and 2) 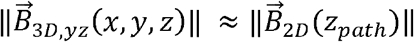. The first criterion was necessary to ensure the magnetic flux lines acting on the MFCs were horizontal (i.e., the only possible y-locations of the tubing were above magnets 1, 3, or 5). The second criterion matched the norm of the magnetic flux in the yz-plane 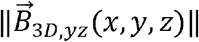 from the 3D simulation to the norm of the magnetic flux 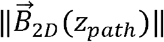 from the 2D simulation in Fig. 2. Satisfying these two criteria allowed the matching of the 3D flux concentrated by the MFCs to what was simulated in 2D, and thus the value of 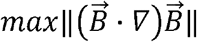 in the tubing cross-section. The output of step 3 was six sets of 3D candidate paths translated from parameter space that recapitulated our target magnetophoretic force profile. We limited our number of potential paths to six, but many potential routes through parameter space, and thus through physical space, could be conceived.

In step 4, the six sets of 3D candidate paths were evaluated using a cost function (Fig. 3D) to select for the path that minimized the following functions: 1) 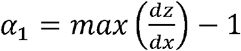, which selects for paths with gradual slopes for the tubing to follow in the xz-plane. This function avoids paths with sharp bends which could lead to difficulties in threading the MFCs wires through the 3D printed part. 2) 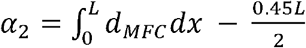, which selects for MFC paths with gradually varying d_MFC_ from 0.45 to 0 mm across the total length L of the path to facilitate the 3D printing and the threading of the MFCs. 3) 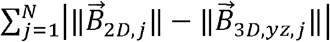, which selects for paths that can best match 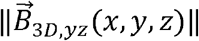 to 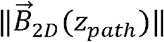 across all N points in the path. The path *θ* with the minimum cost function 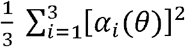 was selected as the final position of the tubing and the MFCs to construct the MSD.

### Standalone MSD and the i-BID achieve high purity and recovery of basophil isolation

To identify the range of flow rates for effective basophil isolation, we first characterized the standalone MSD using enriched WBCs tagged with MNP off-chip. Fig. 4A shows a representative dot plot of our flow cytometry data used to determine purity and recovery of isolated basophils. From 100 μL of whole blood, we expect ~10^5^ non-basophils to be captured on the inner walls of the tubing having a total surface area of ~365 mm^2^ in the MSD. Fig. 4B shows that the purity of isolated basophils was above ~70% for all flow rates tested, with a maximum mean purity of 94.8% (range 90.0 - 96.8%) at 3 mL/hr (n = 5 runs). From 3 to 12 mL/hr, the purity decreased only gently and remained above 80%. The decrease in purity at high flow rates was expected, due to the increased drag forces from the convective flow overcoming magnetophoretic capture forces.

**Fig. 4.**
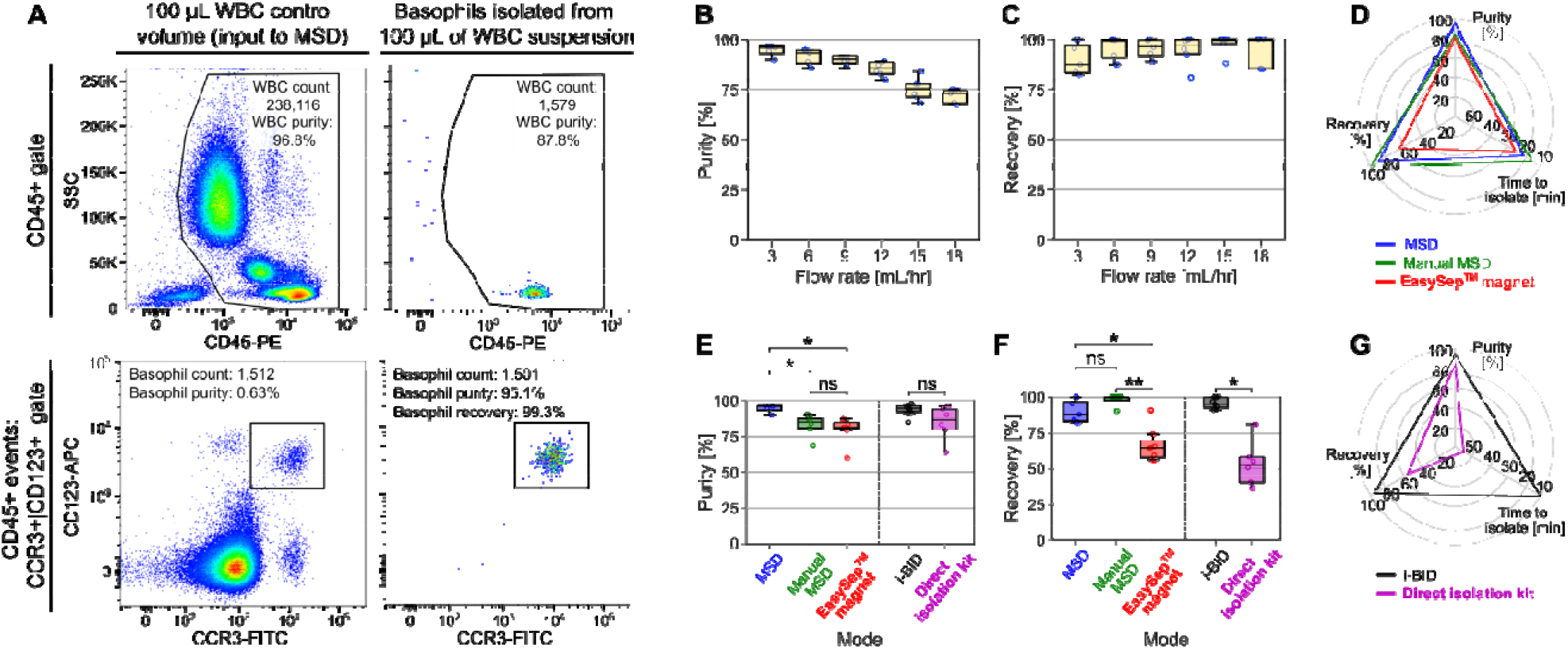
Purity and recovery results. (**A**) Representative flow cytometer data for an experiment characterizing the standalone MSD operated at 6 mL/hr. The dot plots show the initial CD45^+^ cells from which basophils were identified by CD123^+^/CCR3^+^. Purity was calculated as the proportion of basophils in the CD45^+^ gate. Recovery was calculated by dividing the basophil count in the isolated suspension (right column) to the basophil count in the WBC control sample (left column). Basophil purity (**B**) and recovery (**C**) in the standalone MSD characterization. Basophil purity (**E**) and recovery (**F**) compared across different modes of basophil isolation. The MSD mode was operated at 3 mL/hr, and the i-BID was operated at blood injection rates of 3-5 mL/hr. Box and whisker plots presented are Tukey’s box plots (see Methods). \**P* < 0.01, \*\**P* < 0.0001. Spider plots compare the purity, recovery, and time to isolate for (**D**) the syringe pump-driven MSD, the manual MSD, and the EasySep^™^ magnet, and for (**G**) the i-BID and the Direct Isolation Kit.

Fig. 4C shows that the recovery of basophils was above ~80% for all flow rates tested, with all flow rates exhibiting at least one instance of 100% recovery. Across all flow rates tested in our experiments (n = 32), the mean recovery was 94.6% (range 81.0-100%). Slightly lower median recovery (81.0%) was observed at 3 mL/hr. We attribute this lower recovery to non-specific capture of basophils caused by magnetic debris build-up on the tubing walls which was exacerbated when the flow was slow with low drag forces.

Given the high purity and recovery of our standalone MSD and the relative insensitivity to the injection flow rates, we went on to test if we could perform basophil isolation without using any pumps by manually injecting the enriched WBC-MNP/NSAb suspensions through the MSD. This operation allows for a direct comparison of the performance of our MSD with the commercial EasySep^™^ magnet in bulk MACS assays and could enable the broader use of our MSD in labs that do not have ready access to syringe pumps and microfluidics. We instructed the participants to aim for a steady withdrawal of 100 μL of WBC-MNP/NSAb suspension through the MSD in 2-3 min, corresponding to an average flow rate of 6-8 mL/hr. Using this manual operation of the MSD, we achieved 82.9% purity (range 68.7-90.5%) and 98.4% recovery (range 90.2-100%) across all participants (n = 6). Using enriched WBCs from the DLD in the EasySep^™^ magnet, we achieved a purity of 80.1% (range 59.8-87.2%) and a recovery of 65.5% (range 54.7-90.7%) (n = 7 runs) (Fig. 4D).

Overall, the syringe pump-driven MSD exhibited significant improvement in purity (*P* = 0.0015) and in recovery (*P* = 0.0010) compared with the EasySep^™^ magnet (Fig. 4, E and F). The manual MSD offered insignificant improvements in purity over the EasySep^™^ magnet (*P* = 0.52), but it showed significantly better recovery than the EasySep^™^ magnet (*P* = 4.60 x 10^-5^). Compared with the manual MSD, the syringe pump-driven MSD gave significantly higher purity (*P* = 0.011) and similar recovery (*P* = 0.066). The variance in purity using the EasySep^™^ magnet, as determined by an F test (see Methods), was significantly higher than the syringe pump-driven MSD (*F_7,4_* = 9.25, *P* = 0.048) while the recovery exhibited similar variances (*F_7,4_* = 2.36, *P* = 0.42). These results indicate the syringe pump-driven MSD achieved more reproducible isolation purity compared with the EasySep^™^ magnet.

Finally, we tested the performance of the fully integrated i-BID. Compared with the standalone MSD, the integrated i-BID allowed for precise control over the incubation time given its integrated mixing and flow-through design. Although the commercial EasySep^™^ protocol suggested an incubation time of 5 min, we incubated for 3 min instead because it gave a basophil purity of >90%, which already exceeded the purity of bulk basophil isolation using the Direct Basophil Isolation Kit^™^ (Fig. 4D). We chose to operate the i-BID at blood injection flow rates of 3-5 mL/hr because the highest purity was obtained at these flow rates in the standalone MSD, and the probability of clogging at the DLD inlet increased with increasing flow rates. Fig. 4, D and E show that the mean purity of the i-BID was 93.9% (range 85.5-98.2%), and the mean recovery was 95.6% (range 91.0-100.0%) (n = 6 runs). Compared with the commercial Direct Basophil Isolation Kit^™^ with a mean purity of 84.6% and recovery of 52.4%, our i-BID gave significantly improved recovery (*P* = 0.0011), with significantly less variance in both purity and recovery (*F_5,8_* = 9.88, *P* = 0.0057, and *F_5,8_* = 20.52, *P* = 0.00045, respectively).

In our i-BID, the entire process from loading whole blood to obtaining isolated basophils was completed in 8-10 min, which was >4.5-fold shorter than the bulk MACS assay which took ~45 min due to multiple centrifugation steps required to remove RBCs. Furthermore, the integration of DLD in the i-BID workflow enabled basophil isolation from small blood volumes with high levels of recovery. The smallest volume of blood that could be reliably processed by the commercial isolation kit was 300 μL, 3x more than that possible with our i-BID.

### The i-BID does not induce activation status of basophils

For our method to be useful for downstream basophil assays, it is critical that the i-BID does not induce any unwanted activation of basophils and that basophils can still be activated by known stimulants (e.g., anti-IgE). Fig. 5 compares the activation status of basophils isolated from our i-BID vs. basophils in whole blood, incubated with RPMI and anti-IgE, respectively. For the whole blood sample, we followed standard protocol for measuring basophil activation (see Methods). Compared with whole blood control, basophils isolated by the i-BID had similar levels of baseline activation when incubated in RPMI for both healthy and allergic donors (n=3 and n=4, respectively). For all samples, the percent of CD63^+^ basophils was less than 5%, well below the typical level of baseline activation published in the literature (*29*, *57*). This result indicates that our i-BID imparted insignificant activation of basophils. The expression of CD203c, another widely used activation marker (*25*, *27*, *28*), was also unaffected by the i-BID at baseline (Fig. S3).

**Fig. 5.**
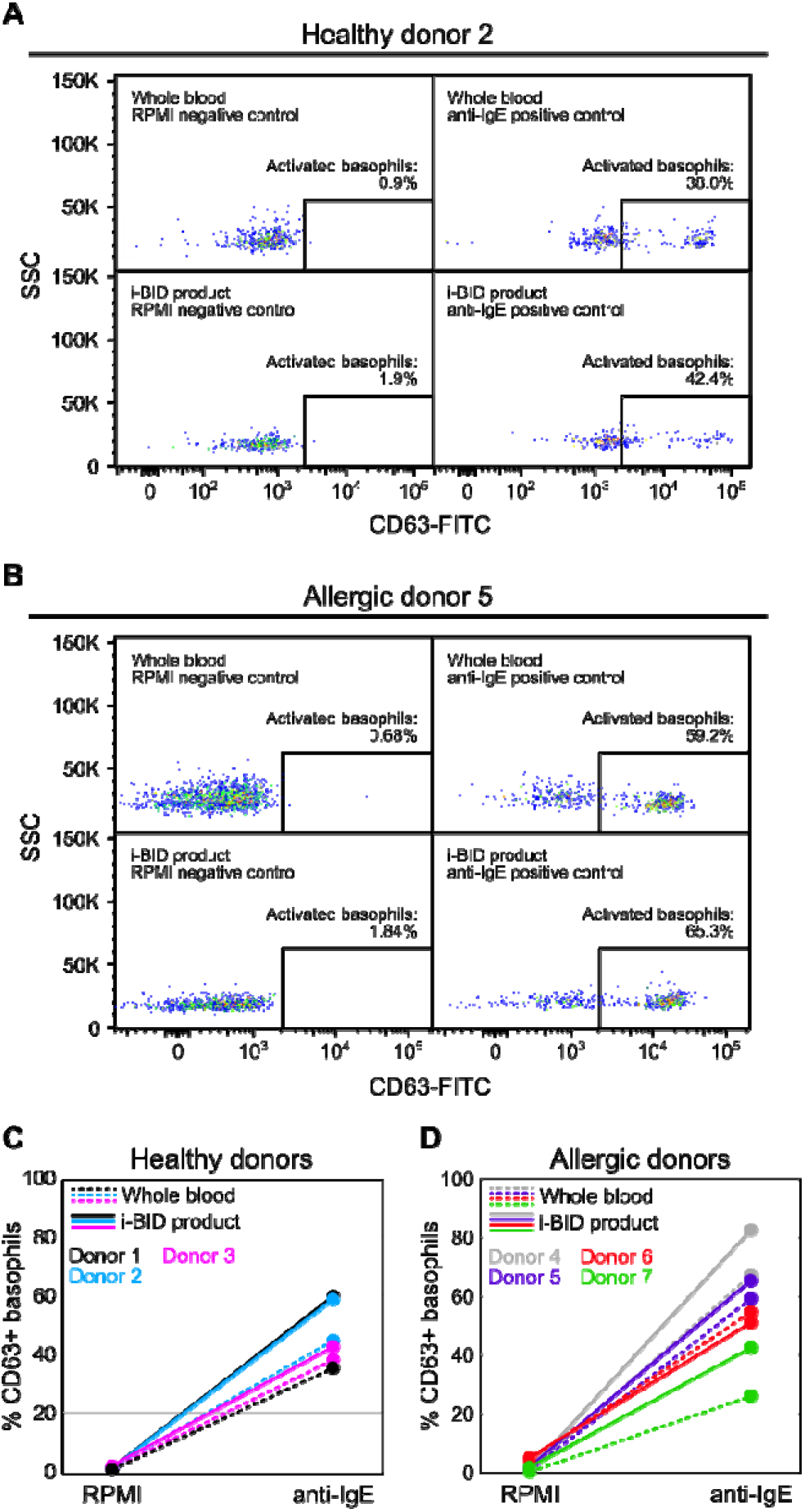
Viable basophils with no artificial activation. The levels of basophil activation (as measured by %CD63^+^) in the i-BID product when subjected to RPMI (negative BAT control) and anti-IgE (positive BAT control) were similar to the levels of basophil activation in whole blood samples that were processed with the standard BAT protocol (*29*). Representative flow cytometer data for (**A**) a healthy donor, and (**B**) an allergic donor, respectively. The levels of basophil activation for (**C**) 3 heathy donors, and (**D**) 4 allergic patients, respectively.

Fig. 5 also shows that basophils isolated by the i-BID retained their ability to become activated in response to anti-IgE. We observed some increase in the level of basophil activation, with a difference in %CD63^+^ ranging from 4.4% (Donor 3) to 23.9% (Donor 1) between the i-BID sample and whole blood control after anti-IgE stimulation. Because baseline basophil activation was not elevated in the i-BID sample, this increase in activation after anti-IgE stimulation may be due to the absence of other WBCs in our enriched basophil sample after isolation by the i-BID. Indeed, some non-basophil WBCs (e.g., eosinophils) present in whole blood, can express FcεRI, a high-affinity IgE receptor (*52*) and bind to anti-IgE that would otherwise interact with and activate basophils. This may have contributed to our observation that basophils in whole blood displayed a lower level of activation compared with the i-BID sample, where enriched basophils were not competing for anti-IgE binding.

## Discussion

With the i-BID, we demonstrated a rapid microfluidic approach (~10 min) for immunomagnetic negative selection of basophils directly from 100 μL of whole blood with high purity (>93%) and recovery (>95%). Compared with the best performing in-bulk Direct Basophil Isolation Kit^™^, the i-BID is >4.5x faster, has higher mean recovery and purity, and its performance is more reproducible with lower variability (Fig. 4, D and G). Rapid isolation of basophils (or other target cells) is important because a lengthy process increases the chance that the native state of the cells is altered. It also increases the likelihood of non-specific uptake of magnetic particles by target cells (e.g., phagocytotic WBCs) through phagocytosis, which can occur on the order of minutes (*53*). Such uptake can lead to the unintended removal of target cells and result in poor recovery. Poor recovery in bulk MACS assays can also arise from sedimentation methods which are often used to remove RBCs and to enrich WBCs. Sedimentation methods have poor leukocyte recovery of < 50% (*54*, *55*) resulting in significant loss of target cells to the RBC-rich sediment. Because of the difficulty in pipetting small volumes of blood samples especially after sedimentation, the practical minimum volume of the blood sample bulk methods can process is typically limited to >300 μL.

Compared with bulk assays, our approach based on microfluidics offers a more rapid and precise way to separate and isolate cells from small sample volumes, due to its ability to control the application of hydrodynamic and magnetic forces precisely to control the path of cells in microchannels. It is also possible to integrate and automate different steps from a conventional assay to minimize the need for manual manipulation and associated variability. While extensive work exists for separating WBC from RBCs (*56*, *57*) and capturing specific cell types (e.g., T cells (*58*) or CTCs (*59*–*67*)) in microfluidic systems, we have only found two reports on microfluidic basophil isolation. Both methods suffered from poor purity (~40-75%) and recovery (~6-64%) (*62*, *63*), far inferior to the performance of immunomagnetic negative selection approaches in bulk and to the performance of our i-BID.

Although some approaches such as those using acoustophoresis were capable of segmenting monocyte, granulocyte, and lymphocyte populations with >83% purity for each subpopulation, they were unable to further distinguish granulocyte subtypes (i.e., basophils, neutrophils, eosinophils) (*64*). Since basophils share similar size, density, and other physical characteristics with other granulocytes (*65*), immunoselection is the most reliable way of isolating basophils. Microfluidic magnetic-activated cell sorting (MACS) is commonly used for on-chip immunoselection (*45*). The primary advantage of MACS is the high specificity with which cell types can be targeted using negatively or positively selecting antibodies. On-chip MACS also allows the implementation of spatially uniform or varying magnetic fields to sort magnetically-tagged cells (*45*, *49*). This capability enables the deflection (*46*) or capture (*47*) of magnetically-tagged cells with varying levels of magnetic susceptibility in regions of different magnetic strengths.

Our i-BID has a few important differences from previous on-chip MACS devices. For example, in one on-chip MACS platform for the immunomagnetic negative selection of CTCs (*46*), two regions with different magnitudes of magnetic flux were used to first deflect highly magnetic cells (>10 magnetic beads per cell) and then deflect lowly magnetic cells (≥1 bead per cell) to waste outlets. While this design demonstrated high purity and recovery of CTCs from a large volume of sample, it required sequential manual steps (i.e., the product from one MACS run was re-labeled at a higher magnetic bead concentration and re-run through the device), multiple off-chip magnetic labeling steps, and was highly dependent on precise alignment of the channels with the external magnetic field.

In contrast to the extreme rarity of CTCs which require large volumes of blood to be processed (~25 mL (*46*, *57*, *66*, *67*)), basophils are more abundant, despite being the rarest granulocyte. We were able to recover >1000 basophils from 100 μL volumes of blood in microfluidic channels with no manual steps. This fact allowed us to rely on DLD channels for WBC enrichment from whole blood. DLD gave high basophil recovery reliably using flow rates that were compatible with the downstream MSD. The low-volume blood requirement also allowed us to operate our i-BID in capture mode instead of deflection mode. The utility of the i-BID centered around a simple, yet effective flow-through MSD with one inlet and one outlet that applies an exponentially increasing magnetic force on magnetically tagged non-basophils to capture them on the inner walls of tubing. With an exponentially increasing magnetic force, non-basophils were removed gradually at the beginning of the MSD without clogging the tubing. A rapidly increasing high-magnitude magnetic force field towards the end ensured that any lowly tagged cells and magnetic debris were removed. With no additional buffer injection required, the MSD is easily integrable with upstream and downstream microfluidic processes. Since a new tubing can be inserted into the MSD without the need to fabricate new microchannels or re-align the flow path with the magnets, our design presents significant advantage over existing microfluidic MACS devices which require microfabrication of ferromagnetic structures and their careful alignment with the microchannel (*45*, *49*).

We have shown that our i-BID did not alter the activation status of basophils for healthy and allergic individuals. The preservation of basophils in their native state is important for downstream analysis such as the basophil activation test and RNA sequencing. However, this part of the study was limited to seven subjects only. Further characterization with more patients is therefore required to validate that our system does not alter basophils.

While our study focused on the isolation of basophils, the methodologies we developed can be applied directly to isolate other subsets of leukocytes (e.g., dendritic cells, neutrophils, progenitor cells), as long as negative selection antibodies for these cell types are available. In addition, our model-driven design pipeline and the use of a 3D-printed scaffold to guide the flow path are expected to be applicable to non-magnetic separation modalities that use other inhomogeneous force fields in 3D (e.g., acoustophoretic, inertial, electrophoretic separation).

We have focused our isolation on a small volume (100 μL) of blood because the ability to rapidly isolate basophils, or other WBC subtypes, from small volumes of unprocessed blood can enable the development of rapid point-of-care clinical diagnostics by removing the need for lab equipment (e.g., flow cytometers and centrifuges) and highly trained personnel. We believe that our i-BID platform, performed on basophils, has strong potential to accelerate immune functional assays for this cell type, such as the basophil activation test for allergy assessment (*24*). It will also be of interest to apply the i-BID platform to isolate other cell types, such as neutrophils to investigate their dysfunction for sepsis diagnosis (*68*), or T-cells for enzyme linked immunosorbent assay (ELISA)-based interferon-gamma (IFN-*γ*) release assays for tuberculosis diagnosis (*69*).

## Materials and Methods

### Blood sample preparation

Blood was collected by intravenous blood draw into heparinized tubes from donors under Institutional Review Boards IRB# 52850. For experiments to characterize purity and recovery, blood samples were kept on ice for at most 3 hours prior to injection into the microfluidic device. To assess basophil activation status, we processed blood samples (kept on ice) within 24 hours of collection.

For experiments characterizing the fully integrated i-BID, we mixed 100 μL of unwashed heparinized whole blood with 1-2 μL of 500 mM EDTA (final EDTA concentration ~5 mM) prior to injection into the i-BID. Heparin served to inhibit free thrombin to reduce platelet aggregation, and EDTA served to chelate calcium to mitigate the formation of clots in the DLD channels (*70*).

For a subset of experiments characterizing the operating conditions of the standalone MSD, we enriched WBCs from 500-600 μL of blood at a time in DLD channels. Because this subset of experiments was not aimed at rapid, fully-integrated basophil isolation with the i-BID, we washed the blood samples to dilute the clotting factors (e.g., fibrinogen and thrombin) and mitigate clogging in the DLD channels. We had observed that injecting whole blood directly into the DLD channels led to considerable build-up of cells at the DLD entrance. The wash was performed by diluting blood ~3000x in 5 mM EDTA in PBS (Ca^-^/Mg^-^), centrifuging the suspension (500 G, 5 min), and aspirating the supernatant for a final volume and hematocrit approximately equivalent to that of the starting whole blood sample.

For all experiments, we added stains to the blood samples to identify basophils in flow cytometry to characterize their purity and recovery after the microfluidic operation. Per 100 μL of blood, we added 3 μL of anti-CD123-APC (clone 7G3, BD Bioscience), 4 μL of anti-CD45-PE (clone HI30, BioLegend), and 4 μL of anti-CD193-PerCP/Cy5.5 (clone 5E8, BioLegend), and incubated the sample for ~18 minutes on ice. Following the stain incubation and prior to injection into the microchannels, we added 22 μL of DLD running buffer (2% FBS, 1 mM EDTA in PBS) for each 100 μL of blood to obtain a final sample volume of 133 μL (blood concentration of ~75% v/v). This step was performed to reduce the chance of clogging.

### Design and fabrication of the deterministic lateral displacement (DLD) device and the mixer

Both the DLD and the mixer were fabricated in poly(dimethylsiloxane) (PDMS) using soft lithography. The master molds were fabricated in SU8-2050 using transparency masks (CAD/Art Services Inc.) and a Quintel Q-4000 mask aligner. The DLD/mixer microfluidic chip comprised two layers of PDMS. The bottom layer contained the DLD channels, and the top layer contained the mixer channel. The two-layer design was necessary for parallelizing the DLD channels as described below.

For the DLD, we adapted the design from Feng *et al*. (*71*). To improve the throughput and to reduce the rate of clogging at the entrance of the DLD arrays, we used four parallel DLD channels. Each DLD channel had a height of 50 μm, a pillar diameter of 22 μm, a gap of 13 μm, a row shift fraction of 1/30, and an approximate critical diameter of ~3.6 μm (see Fig. 1B) (*71*–*73*). We coated the DLD channels with Pluronic F68 (3% w/v, Alfa Aesar J66087 Poloxamer 188) to reduce the incidence of cell adhesion and the likelihood of clogging (*74*). Pluronic F68 was left in the channels for at least one hour, after which it was thoroughly flushed with the DLD running buffer (2% FBS, 1 mM EDTA in PBS) prior to use. Basophil recovery from the DLD was >99% for all blood injection flow rates tested (2-6 mL/hr) when injecting blood that was washed with 5 mM EDTA in PBS (Ca^-^/Mg^-^) (Fig. S1).

After the DLD, MNP and NSAb were introduced and mixed with the WBCs from the DLD using a mixer consisting of a Herringbone-grooved serpentine channel (*48*). The MNP and NSAb were obtained from the EasySep^™^ Human Basophil Isolation Kit supplied by STEMCELL (catalog #17969). The NSAb mixture negatively selected for basophils with a mixture of antibodies targeting non-basophil surface antigens (e.g., CD2, CD3, CD14, CD15, CD16, CD19, CD24, CD34, CD36, CD45RA, CD56, glycophorin A) (*75*). The mixer channels were 200 μm wide and 70 μm tall with groove heights of 30 μm. In our experiments, the flow running through the device was between 3-18 mL/hr, corresponding to a Reynolds number between ~6 to 37, well within the mixer design’s operating conditions (*48*). MNP and NSAb reagents were injected at a rate necessary to maintain a target MNP/NSAb:WBC ratio of 50 μL:1 mL. Their injection rate was thus a function of the expected flow rate of the enriched WBC suspension exiting the DLD channels. For example, to achieve this target ratio for a WBC suspension exiting the DLD stage at 3 mL/hr, we injected MNP/NSAb at 0.15 mL/hr. After use, we flushed the channels with 10% bleach and deionized water. If all debris was removed, the channels were re-coated with F68 and reused.

### Fabrication of the magnetic separation device (MSD)

The Halbach array consisted of five N52 bar magnets (1.5” by 0.75” by 0.75”, Super Magnet Man, catalog #Rect1526). We used a stereolithography 3D printer (Form2, Formlabs) to print the MSD scaffold in Grey Pro resin with a layer resolution of 50 μm. The scaffold consisted of a slot to secure the Halbach array, passage tunnels to position a fluidic tubing (medical grade polyethylene tubing, 762 μm inner diameter, Scientific Commodities, catalog #BB31695) and steel wire MFCs (low carbon 1008 steel, 0.029” dia., McMaster, catalog #8870K15), and openings along the passages to facilitate the removal of uncured resin (see Fig. S4). We chose a tubing with an inner diameter of 762 μm because its cross-sectional area and the corresponding internal surface area were sufficiently large to retain captured non-basophils and unbound MNP without obstructing the flow of basophils, while it was sufficiently small so that the magnetic flux from the MFCs could reach the center axis of the tubing.

We chose to have passes 1 and 4 traverse over magnet 3 (Fig. 1C) because the far-reaching magnetic field over magnet 3 could accommodate low magnetic forces in pass 1 and high magnetic forces in pass 4. Despite the adjacency of passes 1 and 4, no interference occurred in the magnetic field due to the proximity of the MFCs in passes 1 and 4. To prevent tubing intersection over magnet 3, it followed that passes 2 and 3 must span magnets 1 and 5, respectively.

### Flow control of the integrated basophil isolation device (i-BID)

In all experiments, we loaded the blood cells and the basophil isolation reagents in medical grade polyethylene tubing, which were connected to syringes filled with DLD running buffer. To prevent the cell suspensions and reagents from back flowing into the syringes, we used air plugs (5-10 mm long) to separate them from the DLD running buffer which filled the rest of the tubing and syringes. While the air plugs could introduce a source of compressibility into the i-BID, their effects were not observed during steady-state operation.

All isolation experiments were performed at room temperature. Blood flow was actuated with a 1 mL syringe (HSW Norm-ject) and MNP/NSAb flows were each actuated with 250 μL glass syringes (SGE Analytical Science) to prevent syringe deformation under pressure. DLD buffer injected into the DLD/mixer and isolated basophils withdrawn from the outlet of the MSD were actuated with 10 mL syringes (BD Biosciences).

We wrote a custom control program in Python to operate the syringe pumps (Chemyx Inc.). The control code facilitated loading the blood sample, ramping up the flow rate to a constant steady state, and stepping through a series of user-defined operation modes (i.e., injecting blood, flushing the DLD channels, and passing the cells through the MSD). A slight latency in writing and running commands to the pumps across USB serial ports was expected and could lead the contamination of WBC product by RBCs. To avoid this lag, we implemented threading in Python to instantiate multiple processes in parallel and update all pumps simultaneously.

Prior to injecting blood, we initiated the injection of MNP/NSAb and the withdrawal from the DLD/mixer outlet while gradually stepping up the DLD buffer flow until steady state was reached. We fixed the DLD buffer flow rate (Q_buffer_) at 5x the flow rate at which the blood sample was injected (Q_blood_) into the DLD. The flow rates were chosen to keep the average velocities the same in all channels feeding into and out of the DLD arrays. We fixed the flow rate of the mixed WBC-MNP/NSAb suspension (Q_out_) by withdrawing from the outlet of the MSD at a rate equal to the sum of the injection rates of MNP, NSAb, and the WBC outlet flow rate (see Fig. 1A). The RBC waste outlet tubing was submerged in the fluid of the waste collection container to avoid pressure fluctuations due to dripping and maintain a constant pressure at the waste outlet. After the entire volume of blood was injected into the DLD/mixer channels and right before the air plug entered the channel, we advanced the Python control program to the next step to halt blood injection, flushed the DLD channels with DLD running buffer, and ramped down the injection of MNP/NSAb as the WBC-MNP/NSAb suspension continued through MSD.

### Characterization of the performance of the standalone MSD

In a subset of experiments to evaluate the effectiveness of the MSD in isolating basophils as a function of flow rate, we enriched WBCs using the same DLD channels that comprised the i-BID. Aliquots of the DLD product (100 μL, with ~2-3×10^5^ WBCs) were mixed with 5 μL of NSAb and 5 μL of MNP in a 5 mL round-bottom tube with a gentle shake. After 5 min of incubation at room temperature, we injected the mixture into the standalone MSD with a syringe pump in withdrawal mode from 3-18 mL/hr (n =5 runs for all flowrates except for 12 and 15 mL/hr where n = 6 runs).

For a subset of experiments, we tested the operation of the standalone MSD by injecting the enriched WBCs manually from a syringe. Briefly, we asked 6 participants to withdraw the WBC-MNP/NSAb suspension from the outlet of the MSD by hand using a 1 mL syringe. Participants were instructed to steadily withdraw the sample through the MSD in 2-3 minutes resulting in approximate flow rates of 6-8 mL/hr

### Comparison with commercial basophil isolation kit

We compared the performance of the MSD and manual MSD to that of the STEMCELL EasySep^™^ immunomagnetic column-free magnet (catalog #18000). We followed the same process detailed above for enriching WBCs from blood and adding MNP/NSAb to 100 μL of WBC suspension. After 5 min of incubation with MNP/NSAb, we added 3 mL of DLD running buffer and placed the tube in the EasySep^™^ magnet. We allowed 5 minutes for the removal of magnetically tagged cells following the EasySep^™^ protocol. After that, with the tube remaining in the EasySep^™^ magnet, the suspension containing purified basophils was poured into a new 5 mL tube. The cell suspension was then centrifuged at 500 G for 5 min to remove the excess buffer and prepare the cells for flow cytometry.

For direct comparison with a commercial isolation kit that involved no microfluidic steps, we selected the STEMCELL Direct Basophil Isolation Kit^™^ (catalog #19667) that can isolate basophils directly from whole blood. For a fair comparison with the kit using optimal conditions, we isolated basophils from a starting volume of 300 μL, near the minimum starting volume specified in the kit’s protocol. The isolation protocol began with adding EDTA to whole blood in a 5 mL round-bottom tube to reach a final EDTA concentration of 3 mM. Next, 15 μL of the direct kit’s NSAb and MNP were added (50 μL per mL of blood) and incubated at room temperature for 5 minutes. The blood was then diluted with 3.7 mL of 1 mM EDTA in PBS and placed in the EasySep^™^ magnet for 5 min. With the tube remaining in the magnet, the suspension was poured into a new 5 mL tube and another 7.5 μL of MNP was added (25 μL per mL of starting volume). Following another 5 min incubation, the tube was placed in the EasySep^™^ magnet for another 5 min. The process of pouring into a new tube, adding MNP, and letting the tube sit in the magnet was repeated once more (3 total incubation in the magnet), and the suspension of isolated basophils was collected. We centrifuged the suspension at 500 Gs for 5 min to concentrate the cells in a 200-300 μL pellet for staining and flow cytometry.

### Characterization of the purity and recovery of basophils

We quantified the purity and recovery of basophils from the product of our device using flow cytometry (BD FACScan, Cytek Biosciences). Purity was quantified by performing flow cytometry on the MSD product to determine the percentage of events that were SSC^low^/CD45^+^/CD123^+^/CCR3^+^ (Fig. S5) Gating was done in FlowJo v10.8. We followed the convention established by the STEMCELL basophil isolation kit protocol in which purity is determined as the percent of basophils within cells expressing the leukocyte common antigen (CD45) (*40*).

For experiments characterizing the performance of the standalone or manual MSD and the usage of the EasySep^™^ magnet, recovery was quantified by dividing the count of basophils in the MSD product by the count of basophils in the control volume of enriched WBC suspension. Enriched WBC suspensions obtained by DLD were split into 5-6 aliquots each with a volume of 100 μL. One of these aliquots was set aside as the control to determine the expected number of basophils in 100 μL of the WBC suspension. In our recovery quantification, we assumed a homogenous distribution of basophils in WBC suspensions.

For experiments characterizing the performance of the fully integrated i-BID, recovery was approximated by dividing the count of basophils produced by the i-BID by the count of basophils in a control sample of whole blood with a volume equivalent to that injected into the i-BID. This control was stained for CD123, CCR3, and CD45, and WBCs were enriched by lysing RBCs. In our recovery quantification, we assumed a homogenous distribution of basophils in whole blood.

In all experiments, we also collected the RBC waste from the DLD channel, lysed the RBCs, and injected any remaining WBCs through the flow cytometer to verify that <1% of the total expected basophil count was lost to the DLD channel waste (Fig. S1).

To quantify the recovery of basophils using the EasySep^™^ Direct Human Basophil Isolation Kit^™^ by STEMCELL (catalog #19667), we used a 100 μL control volume of whole blood to count the expected number of basophils and multiplied the basophil count by 3 to compare to the direct kit’s recovered count from 300 μL of blood. For the direct isolation kit, we started with 300 μL of whole blood based on our experience with getting better recovery starting with 300 μL compared to 100 μL.

### Characterization of basophil activation status

We performed experiments to examine whether the isolated basophils exhibited any unintended activation in our on-chip isolation process, and whether they could still undergo activation in response to a stimulus. For these experiments, the surface marker stains were not added until after isolated basophils were recovered and subjected to stimulus. Basophils, when recovered, were suspended in DLD running buffer which contained EDTA. Because EDTA chelates calcium ions and basophil activation is a calcium and magnesium dependent process (29), we resuspended the basophils in RPMI with 1 μM of CaCl_2(aq)_ and MgCl_2(aq)_ prior to subjecting them to any stimulus. Following this step, ~100 μL of basophil suspensions were mixed with 100 μL of RPMI or anti-IgE (2 μg/mL) dissolved in 100 μL of RPMI. After incubation for 30 minutes at 37°C in 5% CO_2_, the activation was halted by adding 1 mL of cold 2.5 mM EDTA in PBS (Ca^-^/Mg^-^) to each sample. The samples were centrifuged at 500 G for 5 min at 4 °C. The supernatant was aspirated, and the pellet was resuspended by vortex mixing. We then added 2 μL per stain per sample for identifying basophils (i.e., anti-CD123-APC and anti-HLA-DR-PE/Cy7 (clone L243, BD Biosciences) and for identifying activated basophils (i.e., anti-CD63-FITC (clone H5C6, BD Biosciences) and anti-CD203c-PE (clone NP4D6, BD Biosciences)). The cells were incubated with the stains for 20 min on ice followed by a wash with 3 mL of a stain buffer (2mM EDTA and 0.5% BSA in PBS). Cells were centrifuged at 500 G for 5 min at 4 °C. The supernatant was aspirated, and the pellet was resuspended in ~200 μL of stain buffer for flow cytometry.

We compared the activation of basophils isolated with the i-BID with that in the whole blood control samples. The control samples were subjected to the same conditions as the samples which passed through the i-BID, i.e., EDTA was added to the blood for a 5 mM final concentration, the control samples were left at room temperature for the duration of the i-BID run, and the sample was resuspended in RPMI with 1 μM of CaCl_2(aq)_ and MgCl_2(aq)_ prior to activation. For measuring the activation status of basophils from the whole blood control sample, we used the same procedure for measuring activation in purified basophil and included an RBC lysis step after the stain. We lysed RBCs by adding 4 mL of 1X RBC lysis buffer (Biolegend, catalog #420301) to the resuspended pellet. We incubated the lysis buffer with the cells for 15-20 minutes in the dark at room temperature, and centrifuged the lysed suspension at 800 G for 10 min at 4 °C. The supernatant was aspirated, and the cells were resuspended in 3 mL of stain buffer before another centrifugation at 500 G for 5 min at 4 °C to collect WBCs for flow cytometry.

### Statistical analysis

To determine the significance of improvements in purity and recovery between the syringe pump-driven MSD, the manual MSD, and the EasySep^™^ magnet, and between the i-BID and the Direct Basophil Isolation Kit^™^, we used unpaired two-tailed t-tests assuming unequal variances. The same type of t-test was applied to show insignificance in CD203c ΔMFI between i-BID and the whole blood control. We used a two-tailed two-sample F-test to determine if there was any significance in the variances found for purity and recovery using each mode of isolation. In our results, we reported the F-value with the degrees of freedom indicated in the subscript. For t-test and F-test comparisons, we chose *α* = 0.05 as the minimum type-1 error rate. Tukey’s box plots were used to report purity and recovery across MSD flowrates (Fig. 4, B and C) and isolation modes (Fig. 4, E and F). The centerlines show the median, the box bounds represent the interquartile ranges, and the whiskers extend out to the maximum and minimum data points, excluding outliers. Plots were produced with Python and statistical analysis was performed in Excel.

## Supporting information

i-BID operation

## Acknowledgments

We would like to acknowledge Stanford Nanofabrication Facility and Stanford Nano Shared Facilities for the equipment used for fabrication. We also want to acknowledge Wyatt Slattery from STEMCELL for his correspondence on the usage of the basophil isolation kits. Research reported in this publication was supported by: Stanford BioX Interdisciplinary Initiatives Seed Grants Program, Sean N. Parker Foundation Fund, Crown Family Fund, National Institute of Allergy and Infectious Diseases of the National Institutes of Health grant R21AI149277, National Institute of Allergy and Infectious Diseases of the National Institutes of Health grant R21EB030643. The content is solely the responsibility of the authors and does not necessarily represent the official views of the National Institutes of Health.

## Author contributions

Conceptualization: NC, AMM, SKYT

Numerical simulation and device design: NC

Experimental design: NC

Experimental work: NC, SK, AMM

Data analysis: NC

Visualization: NC

Supervision: SKYT, SJG, KCN

Writing—original draft: NC, SKYT

Writing—review & editing: NC, SK, SKYT, SJG, KCN

Clinical oversight: KCN

## Data and materials availability

The data supporting this publication will be available at ImmPort (https://www.immport.org). Our MSD designs are available on GrabCAD (https://grabcad.com/library/magnetic-separation-device-1). All other data are available upon request.

