## Supplementary Material for "Exponential magnetophoretic gradient for the direct isolation of basophils from whole blood in a microfluidic system"

**This PDF file includes:**

Fig. S1: Representative data showing minimal basophils lost to the DLD channels’ waste.

Fig. S2: Variation in the mean deflection angle across the parameter space and the tubing path.

Fig. S3: CD203c expression on allergic donor’s basophils was unaffected by the i-BID.

Fig. S4: Computer-aided design (CAD) models of two versions of the magnetic separation device (MSD).

Fig. S5: The gating process used for evaluating purity and recovery.

Table S1: Details of COMSOL simulation.

**Other Supplementary Materials for this manuscript include the following:**

Movie S1 (.mov format): Demonstration of the i-BID operation with annotations.

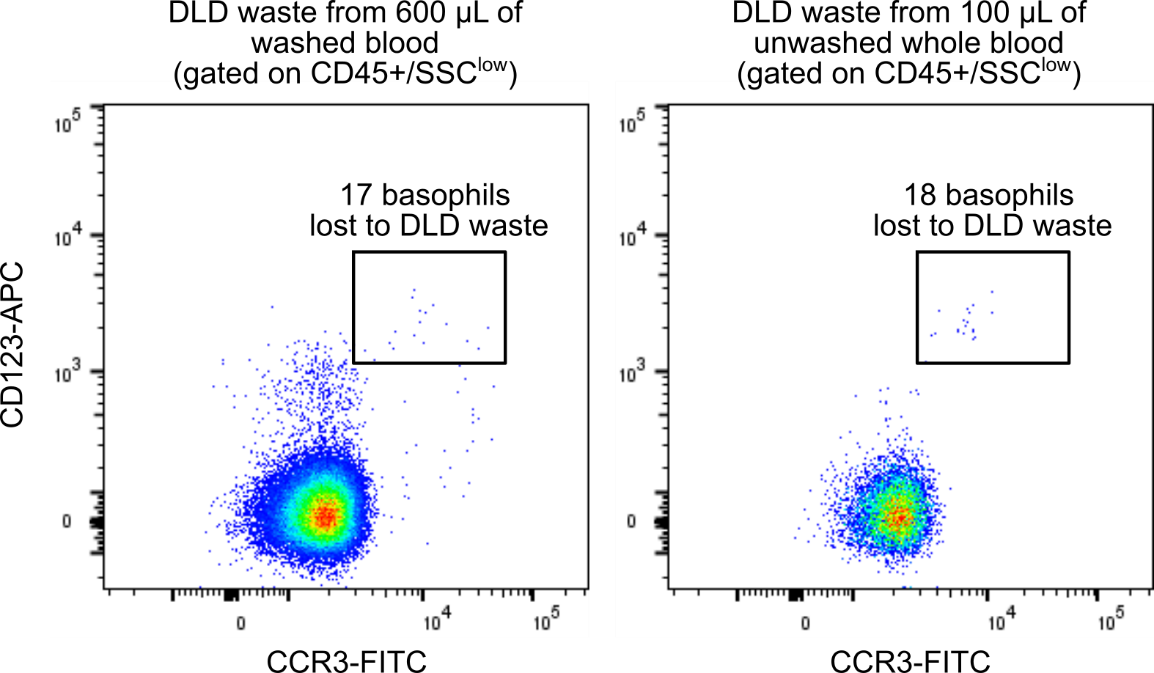

**Fig. S1. Representative data showing minimal basophils lost to the DLD channels’ waste.**

Left: The performance of the deterministic lateral displacement (DLD) channels using blood that was pre-washed in 5 mM EDTA in PBS to dilute clotting factors. This method of blood preparation, used for characterizing standalone magnetic separation device (MSD) runs, allowed the DLD channels to process blood volumes as high as 600 μL with no clogging in the channel and minimal loss of basophils. Right: Unwashed whole blood was more easily affected by clogging at the entrance of the DLD array, but over the course of 100 μL of blood volume, minimal basophil losses were observed. In both cases, <1% of total expected basophils were lost to the DLD channels.

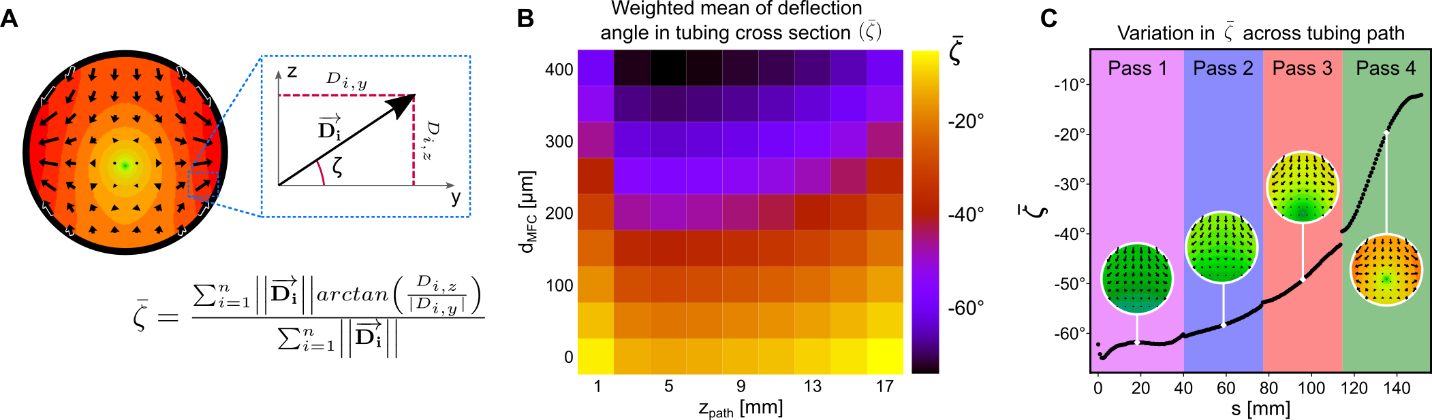

**Fig. S2. Variation in the mean deflection angle across the parameter space and the tubing path.**

(A) All deflection angles of *n* vectors in the tubing cross-section domain are averaged and weighed by their deflection magnitude to give a weighted mean angle, $\overline{\zeta}$. Angles greater than -45˚ (closer to 0˚) indicate a horizontally dominated deflection, i.e., along the global z-axis, that is influenced by the magnetic flux concentrators (MFCs) more than by the Halbach array. (B) Heatmap illustrating the range of horizontally deflected (yellow) to vertically deflected (black) configurations as a function of z_path_ and d_MFC_. (C) $\overline{\zeta}$ approximated along the tubing path. We used the known target maximum magnetic force field value and d_MFC_ at position *s* to look up z_path_, and we used z_path_ and d_MFC_ to look up the corresponding $\overline{\zeta}$ from a smooth cubic interpolation of the heatmap in B. The inflection points are attributed to changes in domain discretization when the values of z_path_ and d_MFC_ are varied and the model is re-meshed. The discontinuities between passes are attributed to edge effects at the ends of the magnets where the tubing turns to run across the Halbach array again.

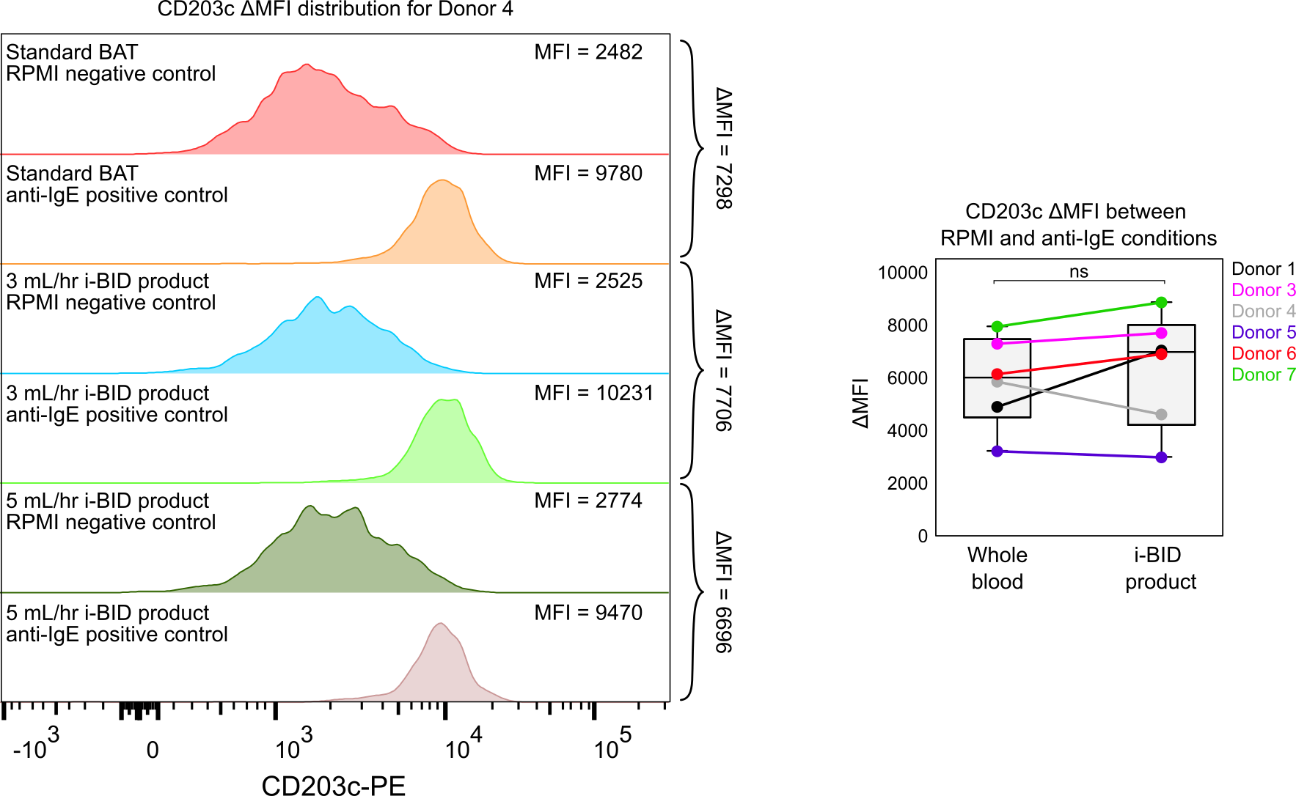

**Fig. S3.** **CD203c expression on allergic donor’s basophils was unaffected by the i-BID.**

CD203c is a commonly used basophil activation marker in addition to CD63. There was an insignificant difference (*P =* 0.69) in CD203c delta mean fluorescence intensity (ΔMFI = MFI_anti-IgE_ - MFI_RPMI_) between i-BID basophils and whole blood control which was processed with a standard basophil activation protocol (see Methods).

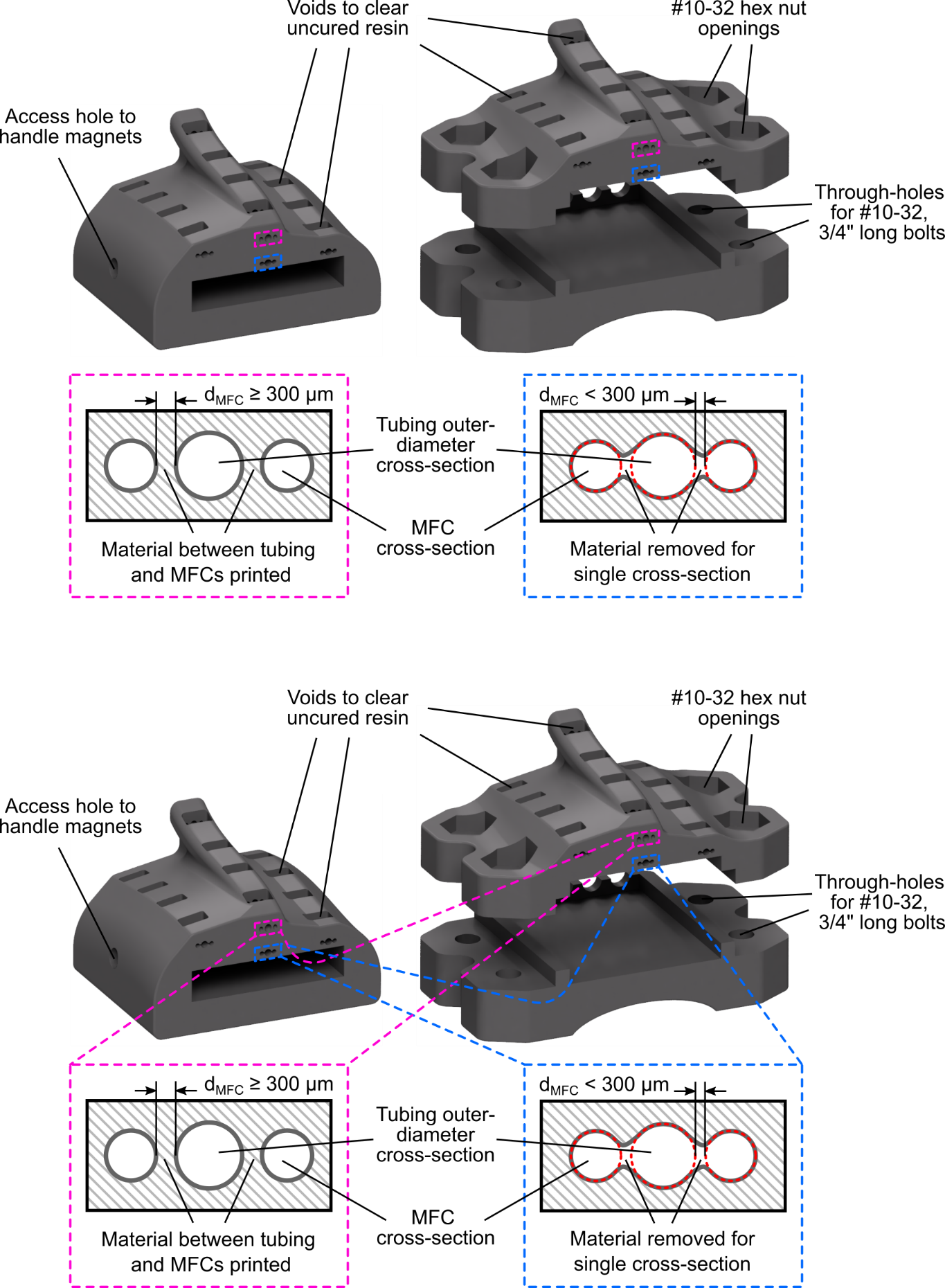

**Fig. S4. Computer-aided design (CAD) models of two versions of the magnetic separation device (MSD).**

We used Formlab’s Grey Pro^TM^ resin for its rigidity. The 3D-printed scaffold surrounded the magnet housing with ample material so that the magnets would not deform the MSD when repelling each other in a Halbach configuration. Both CAD models were used in this work interchangeably exhibiting no difference in performance. They both produced identical magnetic force field profiles with the same tubing path and relative positions between the MFCs and the magnets. The tolerances on the single-piece version (left) allowed for a snug fit around the magnets. The two-piece version (right) ensured the Halbach array could be securely sandwiched using aluminum bolts and nuts, and it facilitated loading and unloading the magnets. For d_MFC_ < 300 μm, to circumvent the minimum feature resolution limit of the Form2 printer that we used, we joined the holes to thread the tubing and MFC. The CAD design is available at: <https://grabcad.com/library/magnetic-separation-device-1>

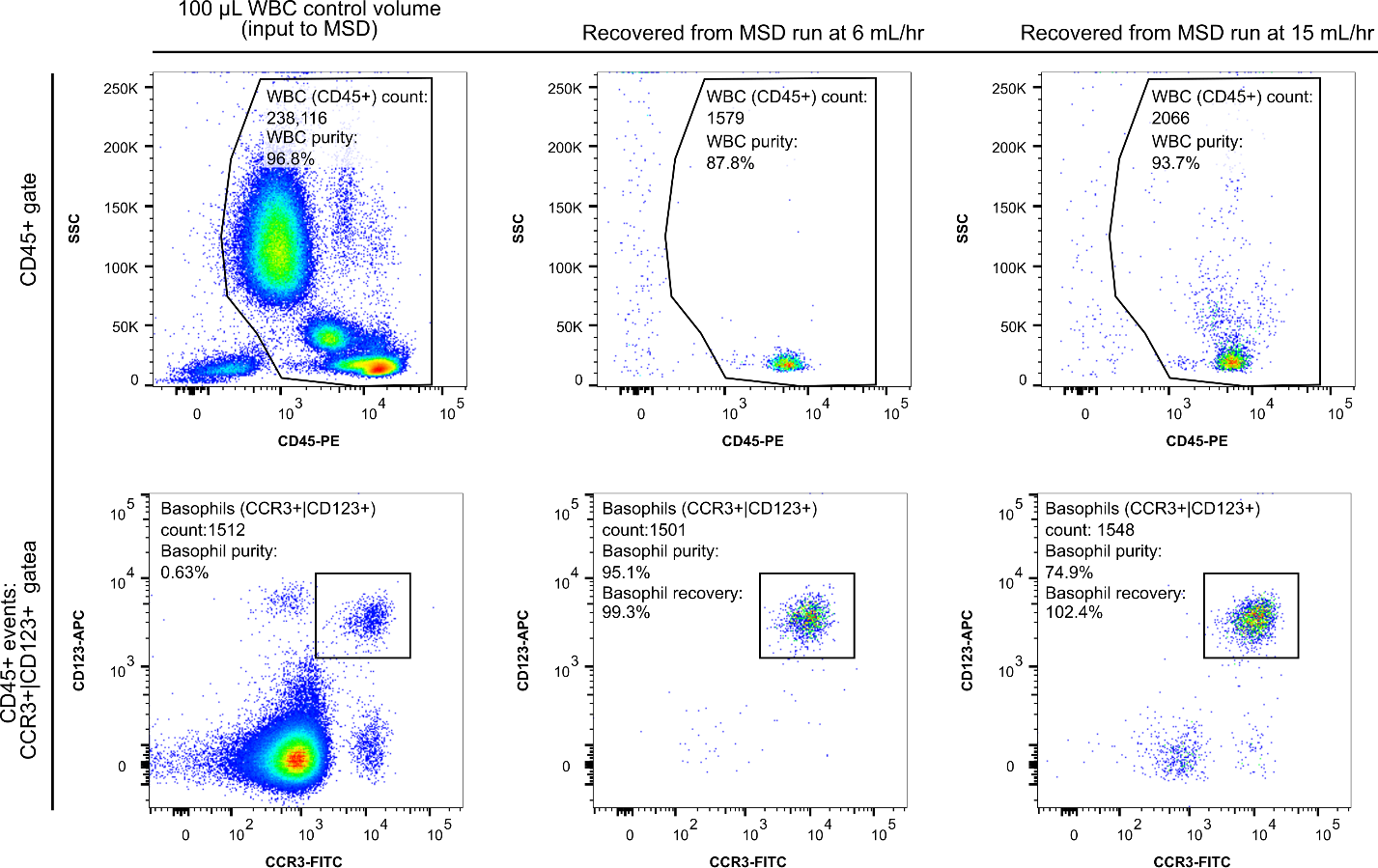

**Fig. S5. The gating process used for evaluating purity and recovery.**

Enriched WBCs from deterministic lateral displacement (DLD) channels were split into 100 µL volumes for testing different flow conditions in the magnetic separation device (MSD). One volume acted as the control to set the CD45 gate and determine the expected number of basophils in the MSD product. Following the convention established by STEMCELL for assessing the purity, basophil purity was determined considering events that were CD45+ only. Events outside this gate in the MSD samples were due to bubbles or debris entering the flow cytometer.

| **Parameters** | |
| --- | --- |
| Tubing outer diameter | 1.22 mm |
| Tubing inner diameter | 762 μm |
| Magnets’ remanent flux density norm | 1.44 T |
| Magnetic flux concentrator (MFC) diameter | 711.2 μm |
| Magnets’ square cross section side length | 6.35 mm |
| Out of plane thickness | 38.1 mm |
| **Materials (all built into COMSOL)** | |
| Low Carbon Steel 1006 | MFCs |
| N52 (Sintered NdFeB) | Magnets |
| Air | All non-magnetic domains (i.e., tubing wall, inner tubing area). |
| **Physics** | |
| Magnetic Fields, No Currents to solve magnetic field | Remnant flux density magnetization model used under Magnetic Flux Conservation domain condition. |
| Coefficient Form PDE to solve magnetic field gradient | Absorption coefficient matrix was cast to an identity matrix. Source Term vector components were defined by the magnetic flux density components. Other coefficient terms were set to zero. |
| **Mesh** | |
| Discretization | Quadratic Lagrange shape functions |
| Mesh resolution | To capture the highly nonlinear magnetic field we partitioned the region near the tubing and MFC for a refined mesh with element sizes from ~2-50 μm. |

**Table. S1. Details of the COMSOL simulation.**

Details of the physical parameters, materials, physics, and mesh are listed above. These values were used for parametric sweeps in 2D to define the parametric space $max\left\| \left( \vec{B}\cdot\nabla\right)\vec{B} \right\|$ and to solve for the 3D magnetic field.

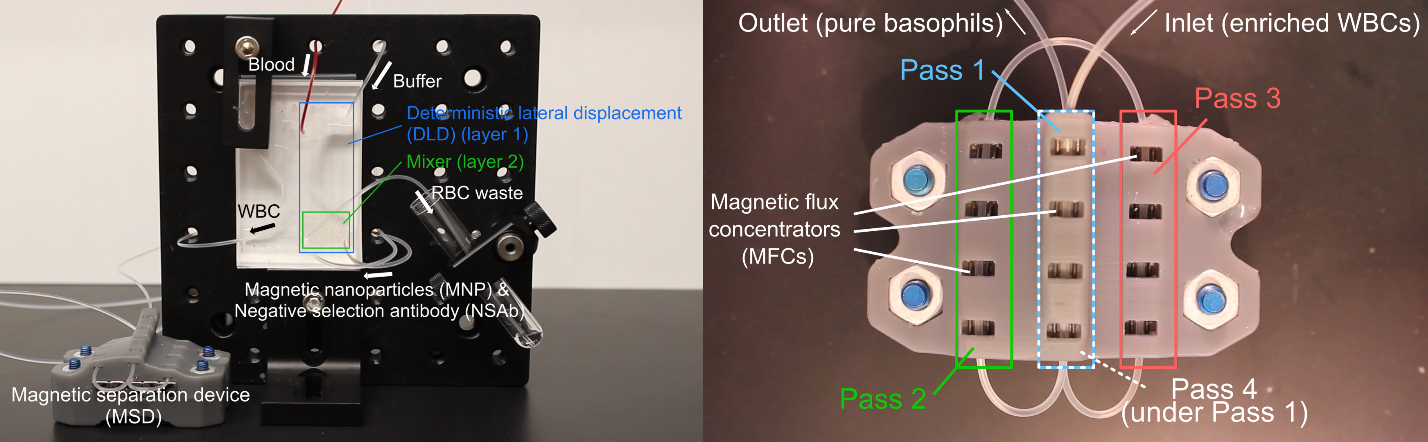

**Movie S1. Demonstration of the i-BID operation with annotations.**

100 µL of whole blood was injected at 5 mL/hr at the deterministic lateral displacement (DLD) channel inlet along with a running buffer and the NSAb and magnetic nanoparticles (MNP) isolation reagents. At 5 mL/hr, with a 3-minute incubation period for NSAb/MNP binding that occurred in a tubing between the poly(dimethylsiloxane) (PDMS) chip and the magnetic separation device (MSD) (not filmed), the isolation took ~8 minutes in total. The air plug that marked the end of the NSAb/MNP-WBC mixture can be seen entering the MSD. The magnetic force in the MSD was strong enough to pull all magnetic material through the air-liquid interface at this air plug, evident by the clear fluid that exits the MSD outlet, compared with the brown-tinted inlet mixture.
